# Predicting Non-Covalent Interactions Between Antioxidants In Biological Membranes Through Molecular Dynamics

**DOI:** 10.1101/2025.02.15.632960

**Authors:** Marving Martin, Benjamin Chantemargue, Gabin Fabre, Johannes Gierschner, Patrick Trouillas

## Abstract

The synergistic association of different polyphenols has gained much interest in the food industry to develop efficient antioxidant cocktails, reducing the concentration of active agents and subsequently potential toxicity. The theoretical description and prediction of such processes is of central interest in this development. This study aims at benchmarking the performance of molecular dynamics (MD) to predict the formation of non-covalent complexes between π-conjugated antioxidants, including two prototypes (quercetin and vitamin E), in a pure 1,2-dipalmitoylphosphatidylcholine (DOPC) lipid bilayer. To reproduce the experimentally observed quenching of vitamin E fluorescence by quercetin, a sphere-of-action quenching model was applied. The predictive capability of MD simulations at capturing this non-covalent association was evaluated with five other π-conjugated potential partners of vitamin E, namely catechin, caffeic acid, myricetin, kaempferol, and galangin. The observed trend agreed with experimental studies, validating again the use of MD simulations as a tool to measure the potential synergism between natural antioxidants.

## Introduction

The survival of an organism strongly depends on its ability to maintain physical and chemical equilibria between inner and outer biological compartments. This is called homeostasis, an optimum state, in which any change is balanced thanks to various biological processes. Disrupting homeostasis creates an imbalance that can quickly lead to physiological disorders. The oxidative stress is one of that kind, describing the imbalance between the production and elimination of reactive oxygen species (ROS) in the cells. Although ROS production is an inevitable consequence of metabolic processes, ROS are also produced by external factors including pollution, smoke, UV light, or more energetic radiations. Although useful for immune cell signaling at low concentrations, an overproduction of ROS induces significant and irreversible damage to cells, disrupting homeostasis. Lipid oxidation (lipoperoxidation), initiated by ROS diffusing into membranes, is a major process responsible for various disorders, mainly associated with modification of membrane permeability and integrity. ^1,2^ To cope with this process, a variety of antioxidants are present in the human body. Their role is to scavenge free radicals or to quench singlet oxygen. Antioxidants can be endogenous like enzymes (*e.g.*, glutathione peroxidase, superoxide dismutase and catalase) and minerals (*e.g.*, zinc, selenium), or exogenous. The main source of exogenous antioxidants is the diet, supplementing the body’s protection against the deleterious effects of ROS. In addition to the antioxidant vitamins (*e.g.*, vitamin E and vitamin C), minerals (selenium), or carotenoids (*e.g.*, beta-carotene or lycopene), polyphenols constitute a broad family of potential natural antioxidants, with a huge chemical variability. They are classified in different molecular sub-classes, including flavonoids, stilbenoids, phenolic acids, and lignans.

Polyphenols are often found in relatively hydrophilic plant compartments. However, their chemical structures made them highly amphiphilic compounds. According to the environment, they can often non-covalently associate in small or large supramolecular arrangements. For example, such a chemical association has long been described to rationalize the stabilization of anthocyanins.^3^ The non-covalent intermolecular association between pigments (anthocyanins) and copigments (*e.g.*, flavonols or phenolic acids) is named copigmentation. This phenomenon stabilizes but also modulates the hue of flowers, vegetables, fruits, or related derivatives like red wine. This intermolecular association is mainly driven by hydrophobic effects, and various (induced) dipolar or quadrupolar effects ^3^ Hydrogen bonding can also be at stake but to a lower extent, mainly because in water, intermolecular interactions between the copigmentation complexes and the solvent molecules may also play a significant role.

This natural process of color stabilization has gained much interest in the cosmetic and agro-food industries.^3,4^ As a side effect, the driving forces of copigmentation has also attracted some interest in the development of antioxidant cocktails. To mix antioxidant polyphenols of different strengths and exhibiting different mechanisms of action is a promising strategy to lower doses of each ingredient and subsequently to lower their toxicity.^5^ To obtain efficient antioxidant cocktails, the activity is expected to be at least additive, unlikely antagonistic^6^, and more likely synergistic.^7,8^ Additive effects are observed when the compounds are either chemically inert to one another or exhibit no mutual interactions, while synergism is obtained when at least two antioxidants interact with each other, possibly inducing the regeneration (re-activation) of one antioxidant by the other.^9^

Various synergistic, antagonistic, and additive effects were summarized by Olszowy-Tomczyk,^9^ based on Oxygen Radical Absorption Capacity (ORAC) and Ferric Radical Absorption Power (FRAP) assays. The knowledge on the effects of antioxidant cocktails against lipoperoxidation is fragmented, although lipids are the major targets of oxidative stress and consequently of antioxidants. In 2012, Skibsted *et al.* used combinations of hydrophilic and lipophilic antioxidants to thermodynamically rationalize the regeneration of the most efficient antioxidants (the most reducing) by less efficient compounds (less reducing).^10^ This study, achieved in liposomes, included vitamin E (α-tocopherol), which was regenerated by plant polyphenols (*e.g.*, quercetin). Synergism may result from adduct formation ^8,11^ or differences in insertion position.^12^ A convincing process to rationalize this synergism assumes the formation of stable intermolecular complexes, allowing the regeneration of antioxidant properties according to their reducing potential.^7,13–15^

Being able to accurately and quantitatively predict the probability of non-covalent interaction between π-conjugated antioxidants is of major importance in supporting the industrial development of efficient antioxidant cocktails. For that reason, computational chemistry methods, in particular based on molecular dynamics (MD), are increasingly explored to provide such predictive studies. However, these methodologies have to be benchmarked against available experimental data. A former joint experimental and computational study in the group investigated synergism of two antioxidant prototypes (quercetin and vitamin E) in a lipid bilayer made of DOPC molecules;^16^ here, the experimentally observed fluorescence quenching was related to the degree of non-covalent association, as computed by MD simulations. One observed that the quercetin-induced quenching of vitamin E fluorescence showed a non-linear Stern-Volmer behavior, which was however not further elaborated in that publication. The theoretical study further allowed to extract bond dissociation enthalpies (BDE) of the most labile hydroxyl group in both vitamin E and quercetin; these were very similar (75.5 and 78.7 kcal·mol^-1^, respectively). This allows H-atom transfer from quercetin to the consumed vitamin E, inducing the regeneration of vitamin E by quercetin; this process is thus likely to occur within these non-covalent complexes.^17–19^

Based on these former results, the current work aims, at one hand, to take a closer look at the fluorescence quenching of vitamin E by quercetin, observed in a DOPC lipid bilayer, discussing different possible mechanisms. On the other hand, we generalize and validate MD simulations as a tool to predict antioxidant synergism for a series of π-conjugated antioxidant partners, targeting five different flavonoids, recognized for their antioxidant properties.

## 1. Experimental and Computational details

### 1.1 Local Concentration of Vitamin E and Polyphenols

The experimental details of vitamin E quenching by quercetin in a DOPC bilayer membrane were described in the publication by Fabre et al;^14^ the Stern-Volmer graph in this previous work was however given *vs*. the quercetin concentration in the suspension, while for the comparison with the simulation, the local quercetin concentration in the membrane is required. The starting point is the molar ratio of vitamin E with respect to DOPC, being from experiment 1:4.^14^ The local concentration of vitamin E is then obtained from the MD calculations (described hereafter), which will allow the extraction of the volume from (i) the area per lipid and (ii) the bilayer thickness, see Section 2.2. This allows to redraw the Stern-Volmer plot as a function of the local concentration (*vide infra*).

### 1.2 Lipid Bilayer

The lipid bilayer system consisted of 72 DOPC molecules solvated in 2880 water molecules and a physiological saline solution (NaCl) of 0.9 g/L. This bilayer system was obtained from CHARMM-GUI membrane builder.^20^ The all-atom MD simulations were performed using GROMACS 2021 ^21^ with the Lipid 17 ^22^ force field to describe DOPC molecules, the TIP3P ^23^ model for water molecules, and the Joung-Cheatham ion parameters.^24^ After energy minimization, the system was heated to 310 K and the semi-isotropic pressure was maintained at 1 bar for 2 ns. Thereafter, a 500 ns production of the system was performed, in which both electrostatic and Lennard-Jones cutoffs were set at 10 Å, while the longer-range interactions were performed using the Particle-Mesh Ewald procedure. The time step integration was set to 2 fs and hydrogen bond constrain was handled by the LINCS algorithm. To ensure the reliability of the molecular organization, the last relaxed structure of the production was used to insert the polyphenols.

### 1.3 Bilayer with Vitamin E and Polyphenols

In a first step, to simulate the experiment of Fabre *et al.*,^14^ we studied the co-insertion of vitamin E in the DOPC bilayer in a molar ratio of 18:72, with an increasing amount of quercetin molecules (4, 7, 9, 11, 14, 18, 25, 29, 36), to match the experimental conditions, were the molecules were likely entirely partitioned (see SI).^14^ In a second step, the combination of vitamin E with six different polyphenols was studied, namely with caffeic acid, catechin, myricetin, quercetin, kaempferol, and galangin, setting an equimolar polyphenol:vitamin E ratio in DOPC, *i.e.*18:18:72. For catechin, a racemic concentration was mimicked (50%:50%, R/S:S/R). The polyphenol:vitamin E pairs were manually uniformly distributed within and between each layer in a pre-stacked configuration below DOPC headgroups, as shown in **Figure 1-a**. This starting configuration was chosen based on previous trials showing a sampling enhancement with this bias (see Figure S1). The molecules were slowly inserted into the bilayer by using an alchemical coupling method, as described by Lundbord *et al*.^25^ The topology of the antioxidants was obtained in their neutral form using the OpenBabel chemical toolbox program.^26^ Their geometries were optimized with B3LYP/cc-pVDZ, using the Gaussian09 ^27^ program, and the partial charges were calculated using RESP ^28^ with the model of Duan *et al*.^29^ As a compatible forcefield, GAFF2 was chosen to define non-bonded and bonded parameters, except the torsional angle between the B and C rings of the flavonoids (quercetin, kaempferol, myricetin, and galangin) that was set following the improved dihedral parameters previously published.^16^ All molecules inserted in the lipid bilayer system were equilibrated as formerly described, using a position constraint on the molecule rings. This allowed each molecule pair to start the MD production stage (1 µs in 3 replicas) in a pre-stacked configuration. AmberTools22 was used to assess area per lipid (APL), order parameter, and z-axis position, and the LiPyphilic^30^ python package was used to assess molecule flip-flops, and orientation angle with respect to the z-axis perpendicular to the membrane surface (see **Figure 1**a). The diffusion coefficients (lag time = 100 ps) was assessed thanks to the GROMACS analysis tool^21^.

**Figure 1.**
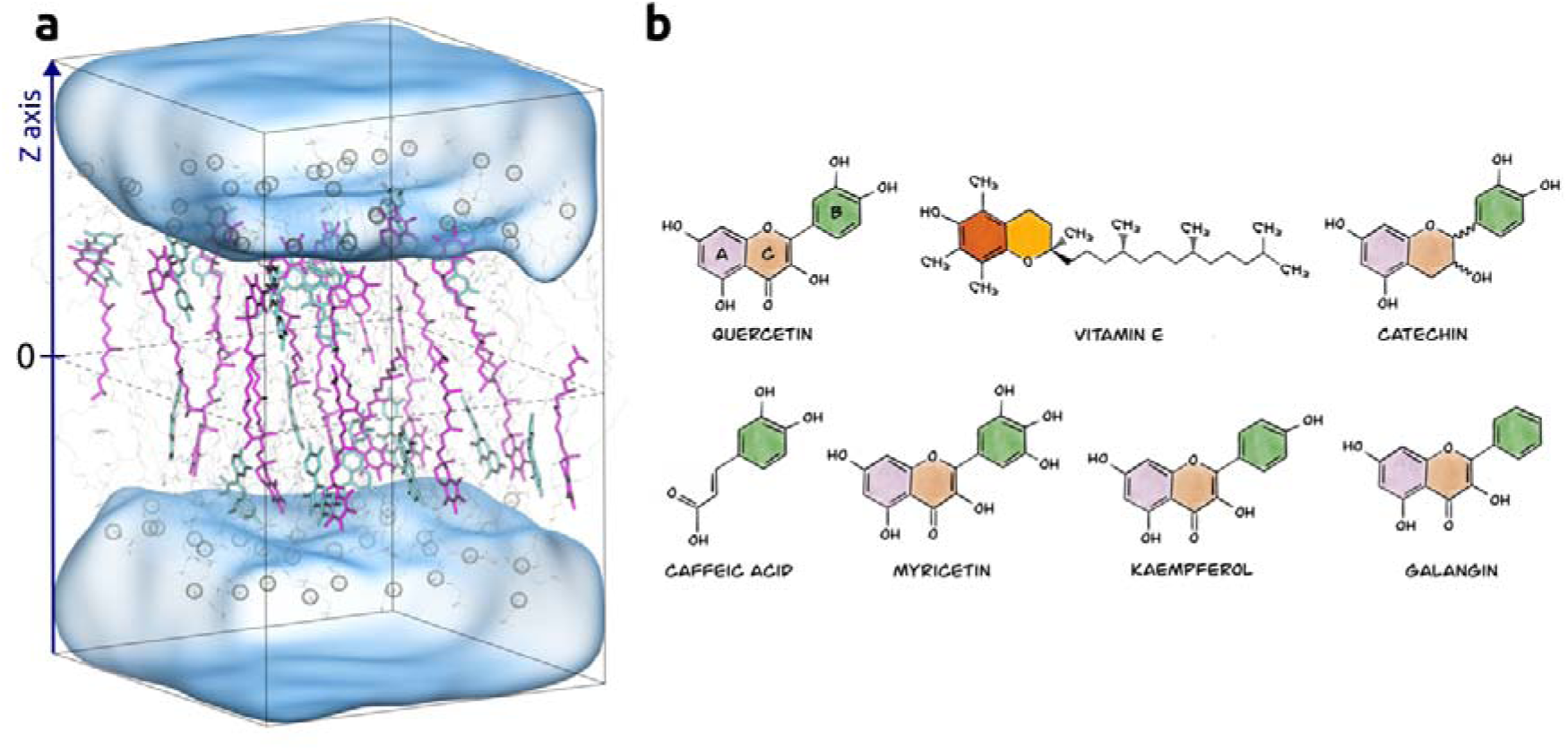
(a) An example of starting configuration with equimolar concentration of quercetin (cyan), vitamin E (purple) in DOPC (18:18:72). DOPC phosphor atoms are highlighted in ochre transparent spheres. Hydrogens have been removed for clarity. (b) List of molecules used in this study with catechin in racemic concentration of R/S and S/R. Flavonoid rings are denoted A, B, and C, in agreement to the regular nomenclature.

## 2 Results and discussion

### 2.1 Structural changes of the bilayer upon lipid-antioxidant interaction

The insertion of polyphenols in phospholipid bilayer membranes is likely to modify the membrane properties in various ways. MD is a suitable technique to monitor such perturbations at an atomic resolution. The impact of vitamin E and quercetin, in a range of concentrations, on different DOPC membrane properties, according to the MD results, is summarized in **Figure 2**.

**Figure 2.**
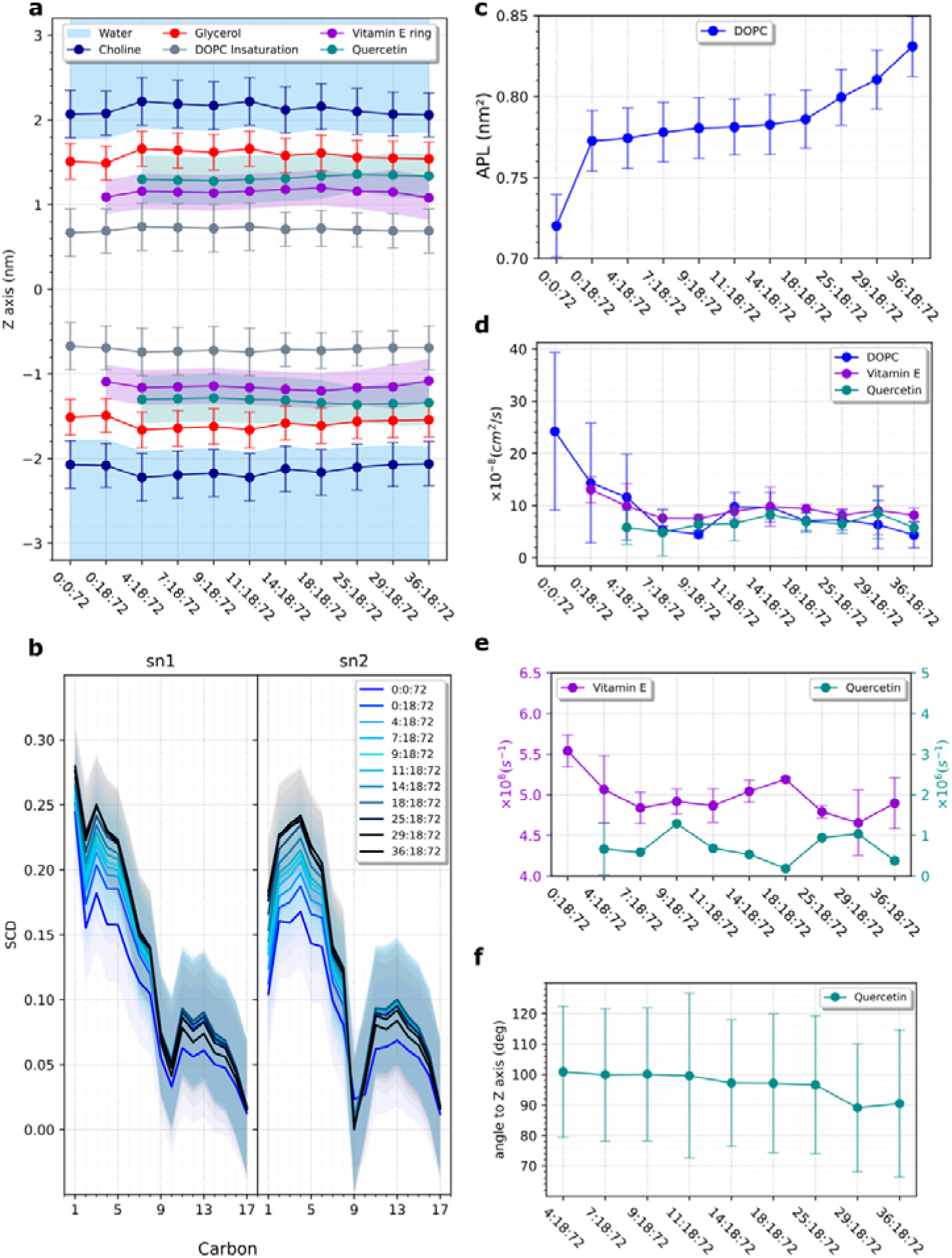
Comparison of membrane properties and molecule behavior between the different quercetin:vitamin E:DOPC systems. (a) Time-averaged position occupied by lipid moieties, polyphenols, and water at 50% of the electron profile density along the Z axis; (b) DOPC acyl chain order parameter; (c) area per lipid, APL; (d) diffusion coefficient of molecules in x,y plane obtain from a linear fit of all mean square displacement; (e) flip flop rate per second ^33^; (f) quercetin angle between membrane normal vector (Z axis) and A-ring-to-B-ring vector.

In the lipid bilayer alone, DOPC molecules diffused in the bilayer at a high diffusion constant, calculated at 2.4 ± 1.5 x 10^-7^ cm^2^/s in 3D space; the system can thus be described as a fluid liquid-disordered phase (*L_d_*), as expected for the given composition, and as confirmed by the order parameter plot (**Figure 2**-**b**). The resulting average values of area per lipid (APL = 0.72 ± 0.02 nm^2^) and Luzzati thickness (D_B_ = 3.6 ± 0.1 nm) agree with the respective experimental data, *i.e.* APL =0.71 ± 0.02 nm^2,31^ and D_B_ =3.6 nm at 303K.^32^ The correct description of these parameters shows the reliability of the MD protocol, based on the adequate forcefield.

Vitamin E is known to be a natural component of cell membranes, which can be provided exogenously (*e.g.*, vegetable oils or nuts). By adding this molecule to the membrane at a 1:4 (Vitamin E:DOPC) molar ratio, we noticed a sharp increase in APL (+0.05 nm^2^), also because of the high vitamin E concentration. Besides, MD simulations also shed light on a drastic drop of DOPC lateral diffusion by about one order of magnitude followed by an increase in the order parameter (**Figure 2**). This phenomenon is mainly explained by the H-bonding interaction between DOPC and vitamin E. As expected, due to its hydrophobic character, vitamin E is located deep in the membrane, with the aromatic rings placed halfway between the DOPC glycerol moieties and DOPC’s unsaturated tail. The overall membrane thickness remained unchanged, partly because of the significant rate of vitamin E flip-flop, assessed during MD simulations in Vitamin E:DOPC system (**Figure 2**-**e**). This mobility along the z-axis goes along with the mobility in the x-y plane, as seen in **Figure 2**-**d**, showing vitamin E with a higher lateral diffusion coefficient than DOPC lipids.

We subsequently studied quercetin co-insertion with vitamin E, and its concentration impact on the bilayer membrane. As explained in the “Experimental and Computational details” section, upon insertion of vitamin E and quercetin, both compounds were engaged in pre-configured π-π stacks. While some pairs remained formed within the first nanoseconds (due to dipole-dipole interactions but also H-bonding), some others quickly split. Then, because of long enough MD simulations to sample the movements of the molecules in the membrane, some pairs were reformed, creating transient non-covalent interactions. By comparing all systems one to each other, the concentration effect was followed (**Figure 2**), highlighting a low- and a high-quercetin concentration groups, where ‘low’ and ‘high’ are defined with respect to the vitamin E concentration (*i.e.*, ‘low’ and ‘high’ mean below or above that concentration, respectively).

#### Low quercetin concentration

The flavonoid is mainly located right under the lipid glycerol moiety. From this position, quercetin can establish π-π stacking interactions with the vitamin E aromatic ring, and mainly H-bonding with water and the DOPC headgroups. The latter interactions with lipid headgroups are predominant for quercetin (> 100%, *i.e.* more than one H-bond per molecule). This is due to the orientation of quercetin, measured as the angle between the A-ring-to-B-ring vector and membrane normal. This angle was about 100° (**Figure 2**-f), in agreement with published data,^34,35^ thus constraining quercetin to align in a relatively parallel fashion with the membrane surface.

#### High quercetin concentration

At equal concentrations of quercetin and vitamin E, the behavior in the membrane changed, making this system a pivot point in the rationalization of concentration effects. Indeed, at this concentration, all involved molecules exhibited a synchronized lateral diffusion (**Figure 2**-d), reaching a plateau of diffusion, as quercetin saturates the membrane. Simultaneously, the two molecules started to move away from each other, and vitamin E inserted deeper inside the membrane at the same position as when it was the only one inserted. This suggests that when quercetin saturates the membrane, the thermodynamic balance of interaction established with vitamin E and the DOPC lipid switches toward another one. A thorough investigation of the MD simulations revealed a significant increase in the size of self-interaction quercetin aggregates, involving up to six members (**Figure 3**). The formation of these aggregates located in between lipid headgroups untangles the rapid increase of the APL (+1nm^2^, see **Figure 2**-c), as well as the shallower penetration of water and reduction of membrane thickness. These results perfectly agree with studies suggesting quercetin crystallization when concentration increases above 24 % molar conc. in DOPC (*i.e.* > 18 molecules for 72 lipid membranes).^34^ At high concentration, the driving forces in these non-covalent aggregates override the H-bonding between quercetin and the lipids, thus constraining quercetin even more parallel to the membrane surface.

**Figure 3.**
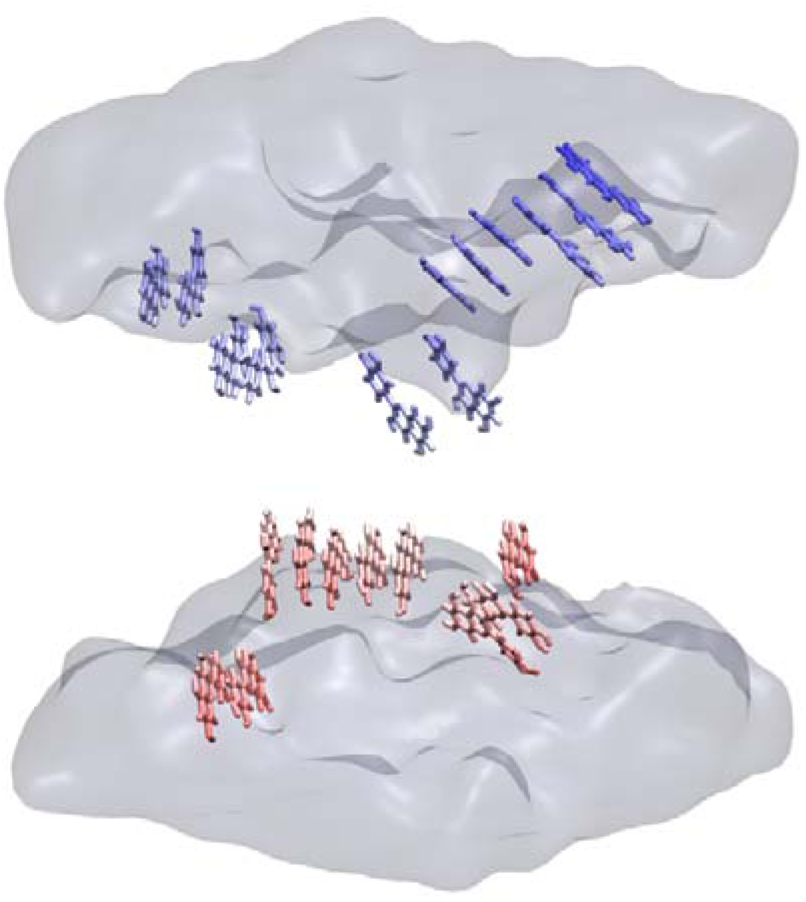
Quercetin saturated quercetin:vitamin E:DOPC (36:18:72) system showing self-interaction quercetin aggregates. The different colors simply differentiate the molecules in each layer of the membrane.

### 2.2 Rationalization of fluorescence quenching

In order to validate that MD can predict the capacity of two antioxidant partners to interact with each other in a lipid bilayer membrane, a thorough understanding of the underlying mechanism of the non-covalent quercetin:vitamin E association is required. To the best of our knowledge, the quenching of vitamin E fluorescence by another antioxidant (quercetin), in a lipid membrane, has been the only implemented experimental technique to monitor this process.^16^

A quantitative evaluation of the data requires the local molar concentration of vitamin E and quercetin, respectively, in the membrane. This can be derived from the MD simulations of the Quercetin:Vitamin E:DOPC system. In the Vitamin E:DOPC (18:72) system, the area of the lipid bilayer is 5.0 nm x 5.0 nm, and the membrane thickness is 5.0 nm. Within this volume, the molar concentration of vitamin E is 0.5 mol/L (see SI for details). Subsequently, the change in the fluorescence quantum yield Φ_F_ (*i.e.* measured by the light intensity under constant excitation conditions and given fluorophore concentration), was plotted against the local quercetin molar concentrations according to the Stern-Volmer (SV) equation:

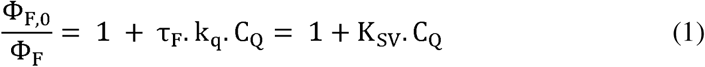

where k_q_ represents the bimolecular quenching constant, *τ*_F_ (s) is the fluorescence lifetime in the absence of a quencher, and K_sv_ (M^-1^) is the SV constant. Φ_F_,_0_ and Φ_F_ refer to the fluorescence quantum yields in the absence and the presence of the quencher, respectively. The SV equation, expressed by quantum yields, is valid for dynamic (collisional) quenching as well as for static quenching,^38^ and should be linear *vs*. C_Q_ (*i.e,* quencher concentration, here quercetin), if only one of the mechanisms is operative.^‡^

As apparent from **Figure 4**, the experiment does not follow the simple linear form of Eq. (1), but it shows a positive deviation. A trivial, but sometimes non-negligible effect, which can cause such deviation, and which is often overseen in literature, is reabsorption (*i.e.* inner filter effect). In the current case, however, considering the overall small concentration of quencher (quercetin) within the suspension, this effect is likely negligible as demonstrated by the I_0_/I value obtained when fitting the experimental data to the adequate reabsorption equation in the SI.

**Figure 4.**
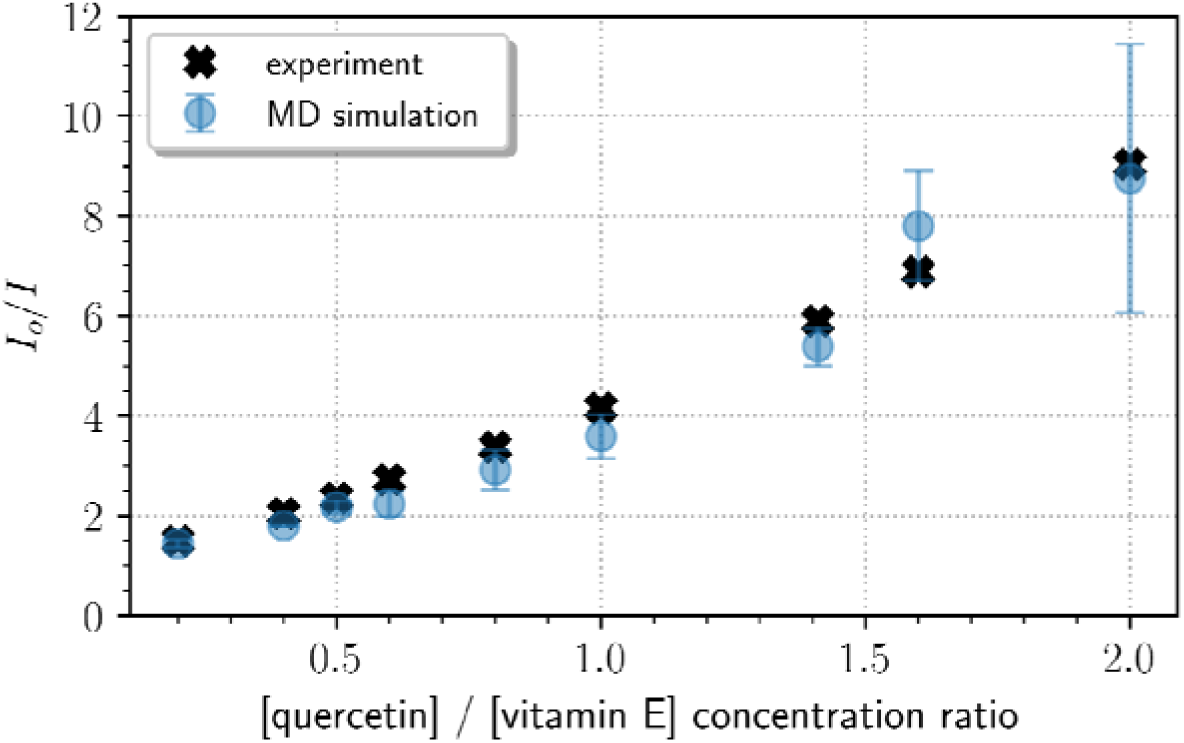
Stern-Volmer plot obtained from MD simulation with a standard deviation (blue) and from Fabre *et al.*^16^ fluorescence experiment (black) of vitamin E quenched by quercetin. Raw data are provided in SI.

Positive deviations from the linear SV relationship are well documented in the literature;^38,42^ empirically, the data can often be fitted by introducing an exponential term in the SV equation:^37^

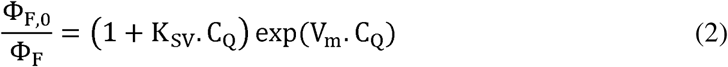

For not too strong deviation from the linear behavior (*i.e.* small V_m_), the exponential function can be expanded into a power series. Truncation after the quadratic term leads to: ^38^

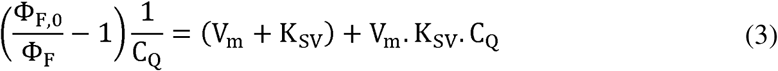

The deviation from the linear SV is commonly interpreted as simultaneous occurrence of dynamic and static quenching, where the latter process is most frequently described as a ‘sphere-of-action’ model, as detailed by Castanho *et. al*.^39–41^

The ‘sphere-of-action’ of a molecule was first introduced by R. Clausius (1858),^42;43^ and was later reconsidered for transient quenching effects,^37^ that occur when both quencher and fluorophore reside within a molar volume Vm. Here, V_m_ defines the action volume occupied by the fluorophore on its random walk during the excited state lifetime τ. As the underlying elementary steps are typically shorter than the fluorescence lifetime,^39^ the quenching can be viewed as instantaneous. Quenching inside the sphere-of-action is enabled by close intermolecular contact, governed by all kind of non-covalent interactions such as H-bonding, dipole-dipole interactions including π-π stacking; such interactions were indeed observed along the MD trajectories.

The quenching curve in **Figure 4**, could be well fitted by the exponential expression of eq. (2) with a high correlation coefficient (R^2^ = 1), giving K_SV_ = 4.42 M^-1^, and V_m_ = 0.51 M^-1^; in simple spherical quenching model this corresponds to an effective radius of 6.^§^

From K_SV_, the quenching constant k_q_ can be estimated from eq. (1), inserting the fluorescence lifetime of vitamin E, τ_F_ ≍1.5 ns,^44^ to give k_q_ = 3·10^9^ M^-1^·s^-1^. In order to interpret k_q_, one may assume, in a first approximation, that dynamic quenching is controlled by a simple Smoluchowski-type diffusion process, expressed by:^45,46^

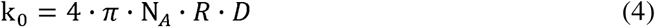

where R = R_F_+R_Q_ is the effective radius of fluorophore and quencher, and D = D_F_+D_Q_ is the diffusion constant. All numbers can be obtained from the MD simulations, giving R_F_ = 2.9 Å (fluorescent aromatic ring) and D_F_ = 1.3 ± 2.5·10^-7^ cm^2^/s for vitamin E, and R_Q_ = 4.2 Å and D_Q_ = 6 ± 3.3·10^-8^ cm^2^/s for quercetin, to give k_0_ = 8.6·10^10^ M^-1^·s^-1^. For such a value of k_0_, the quenching efficiency (*f_q_*) given in eq. (5) would require to be negligible, precluding diffusion-controlled reaction.

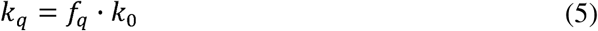

This could also demonstrate, like in other cases,^38^ that eq. (4) is not applicable to membranes. Besides the predominant quenching dynamic mechanism (expressed by K_SV_), static quenching (expressed by V_m_) is significant, giving a weight of x_stat_ = V_m_/(V_m_ + K_SV_) = 0.10. In order to gain further insight into the mechanism, the ratio of quenched fluorescence can also be retrieved from µs-time scaled MD simulations by using equation (6):

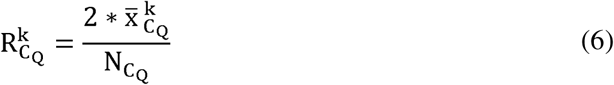

where k is the number of meeting-events between fluorophore and quencher; 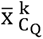 is the number of frames where a meeting-event is observed (*i.e.* minimum distance), averaged over the total number of frames; N_C_Q__/2 is the number of molecule-pairs (*i.e.*, total number of fluorophore and quencher molecules at a concentration C_Q_ divided by 2).

For quercetin and vitamin E, a radius of 7 Å was considered for the sphere-of-action, which corresponds to the sum of the radii of quercetin and the fluorescent moiety of vitamin E, namely 4.2 Å and 2.9 Å, respectively. The latter two values were obtained by considering the two molecules as rigid spheres of which the surface equals their respective trajectory-averaged van der Waals surface (*i.e.*, 221 ± 1 Å^2^, 104 ± 0.6 Å^2^). This value of the sphere-of-action radius, issued from the MD simulation, agrees very well with that obtained above with equation (2) and the curve in **Figure 4**.

Assuming that quenching is ruled by a sphere-of-action mechanism, the ratio equals the loss of fluorescence intensity as defined in equation (7):

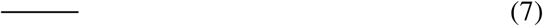

This allows to obtain a simulated SV plot point by point, using equation (8), from each nine [quercetin]/[vitamin] ratios that were simulated by MD (**Figure 4**):

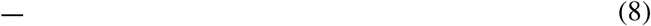

The simulated data presented in **Figure 4** seem to almost perfectly reproduce the experimental data with a mean absolute error (MAE) of 0.41. This strongly supports the probability of a sphere-of-action quenching and also demonstrates that MD simulations can reproduce the overall driving forces that are responsible for the transient interactions ruling this mechanism. We can therefore assume diffusion of two molecules until they collide at a distance of *ca.* 7 Å. In any case, albeit highly promising, we note the limitation of the MD at large number of quenchers (quercetin); in fact, the standard deviation increases, particularly for [quercetin]/[vitamin E] ratio higher than 1.5. From this point, the discrepancy between MD and experimental data could be rationalized by the need for greater sampling time at high concentrations, where the molecular system is likely to accumulate in local minima due to an increasing size of quercetin aggregates.

Finally, we stress that the elucidation of the exact mechanism for the quenching process within the sphere-of-action model remains intricate. Besides trivial quenching by reabsorption (which we could exclude; *vide supra*), quenching could occur *via* photoinduced electron transfer, or (Coulomb-type) energy transfer (Förster transfer);^36^ this is even complicated due to the high quencher concentrations in the current case, which may require more complex multi-step considerations.^47,48^ Clear distinction can only be made by more sophisticated experimental studies (in particular transient absorption and fluorescence experiments), which are however not available to date on these systems.

### 2.3 Quantification of non-covalent association

Non-covalent complex formation between the extended π-conjugated compounds is characterized mainly by close π-π stacking (face-to-face; driven by dipole-π and π-π interactions),^36,49^ or by T-shaped (edge-to-face) arrangements (driven by CH-π-interactions),^3^ The stacking probability (*i.e.* of the interaction between quercetin and vitamin E) can be directly accessed by MD simulation. The pairs are considered as stacked arrangements, when they meet the following two conditions: i) the distance between the two centers of geometry of the aromatic rings is below or equal to 5 Å for face-to-face and 5.5 Å for edge-to-face arrangements; ii) the interplanar ring angle is either 0° ± 5° for parallel stacks or 90° ± 5° for T-shaped conformations. This condition was thoroughly discussed in the literature and it was chosen based on multiple articles,^50–54^ to account for all quick re-stacking situations. The percentages are presented in **Figure 5**-a.

**Figure 5.**
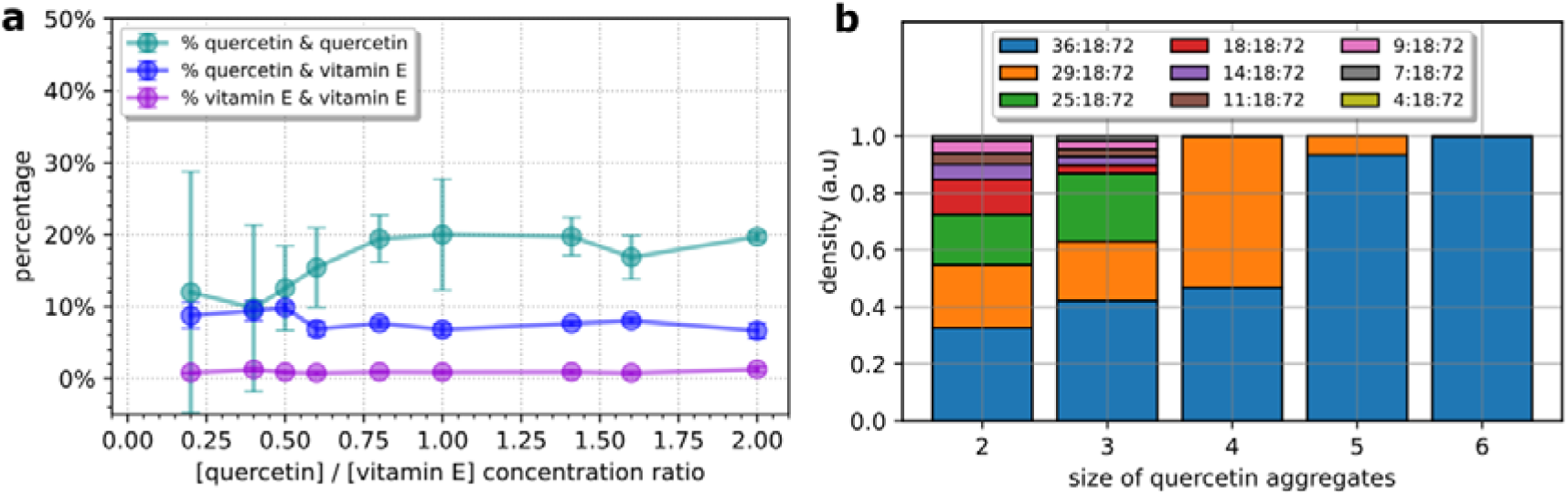
(a) Percentage of stacking interaction in MD simulation by molecule pair embedded in DOPC bilayer membrane. (b) Normalized size distribution of π-π stacked quercetin aggregates in quercetin:vitamin E:DOPC systems.

This graph shows that stacking of vitamin E and quercetin in MD contributes for an average of 8.2 ± 1 % to all meeting events, which agrees with x_stat_ = 10%. On the other side, vitamin E self-interaction is very unlikely (0.9 ± 0.2 %). As seen in the first section, vitamin E is highly diffusive in the membrane and the subsequent amount of flip-flop reduces the possibility of self-interaction. By contrast, as quercetin concentration rises, the amount of π-π stacked self-interaction aggregates increases, before reaching a plateau at around 18.3 ± 1.7 %. This could be explained by better sampling (lower standard deviation) when the quercetin concentration is high, while the molecules are maintained in π-π stacked aggregates as concentration exceed equal molar conditions (**Figure 5**-b), which corresponds to 17 mol%. This agrees with SAXS experiments bringing to light quercetin crystallization above 24 mol% in a similar lipid bilayer system;^34^ here, no quercetin and vitamin E sandwich-aggregates of π-π stacking was observed. This can be rationalized based on the geometrical features of both molecules. To form π-π-stacks, quercetin offers two sides, which easily favor the formation of bigger sandwich-aggregates when concentration rises. Conversely, the methyl group on the asymmetric carbon atom of vitamin E is too bulky to allow π-π stacking on both sides of the molecule, thus preventing the formation of sandwich-aggregates.

### 2.4 Polyphenol structure-activity relationship

To extrapolate the use of MD simulations at predicting the capacity of two π-conjugated compounds to interact with each other (and to rationalize the modes of interaction) in lipid bilayers, we simulated five other equimolar combination with five polyphenols (catechin, caffeic acid, myricetin, kaempferol and galangin) and vitamin E, the simulation systems being named CAT:VIE:DOPC, CAF:VIE:DOPC, MYR:VIE:DOPC, KAE:VIE:DOPC and GAL:VIE:DOPC, respectively, with the following molecular ratio: 18:18:72-.

The different polyphenols revealed different depths of insertion in the DOPC lipid bilayer (**Figure 6**-**a**). Catechin is found to be the less buried in the membrane, followed by caffeic acid, myricetin, quercetin, kaempferol and finally galangin. This time-averaged position along the z-axis follows the octanol/water partition coefficient (XLogP3) assessed on SwissADME website^55^ for each molecule, as XlogP3 equals 0.36, 1.15, 1.18, 1.54, 1.90 and 2.25, respectively. Because of concomitant presence of both the polyphenol and vitamin E and possible interactions between them, the more lipophilic the polyphenol, the less vitamin E sinks deeply in the bilayer. Indeed, as vitamin E is driven closer to the polyphenol, it is gradually pulled away from lipid glycerol moieties. Galangin and catechin are the two extreme cases of the series of the six polyphenols, on the lipophilicity scale. Conversely to catechin, galangin locates at the same depth than the aromatic ring of vitamin E. This favors more contacts and intermolecular interactions inside their sphere-of-action. Both galangin:vitamin E and quercetin:vitamin E couples exhibited the highest probability of π-π stacking interactions (8.9 ± 0.7%; 8.7 ± 0.9%, respectively, as seen **Figure 6**-**b**). Despite being more lipophilic than quercetin, kaempferol appears as an exception as it exhibited a lower percentage of π-π stacking interaction (8.8 ± 0.9%, and 5.4 ± 0.5 % for quercetin and kaempferol, respectively). It must be stressed that H-bonding between polyphenols and DOPC glycerol moieties also contributes to the global balance, which however decreases when the polyphenol is more buried (**Figure 6**-**c**). This observation helps to discriminate comparable percentage of non-covalent interaction as between kaempferol and myricetin. In fact, the latter two share a similar percentage of π-π stacking around 5% (respectively 5.4 ± 0.5% and 5.7 ± 0.4%). However, kaempferol H-bonding interaction with glycerol appears to be less important in the system as compared to the π-π stacking contribution conversely to myricetin. This likely suggests a greater antioxidant action of kaempferol.

**Figure 6.**
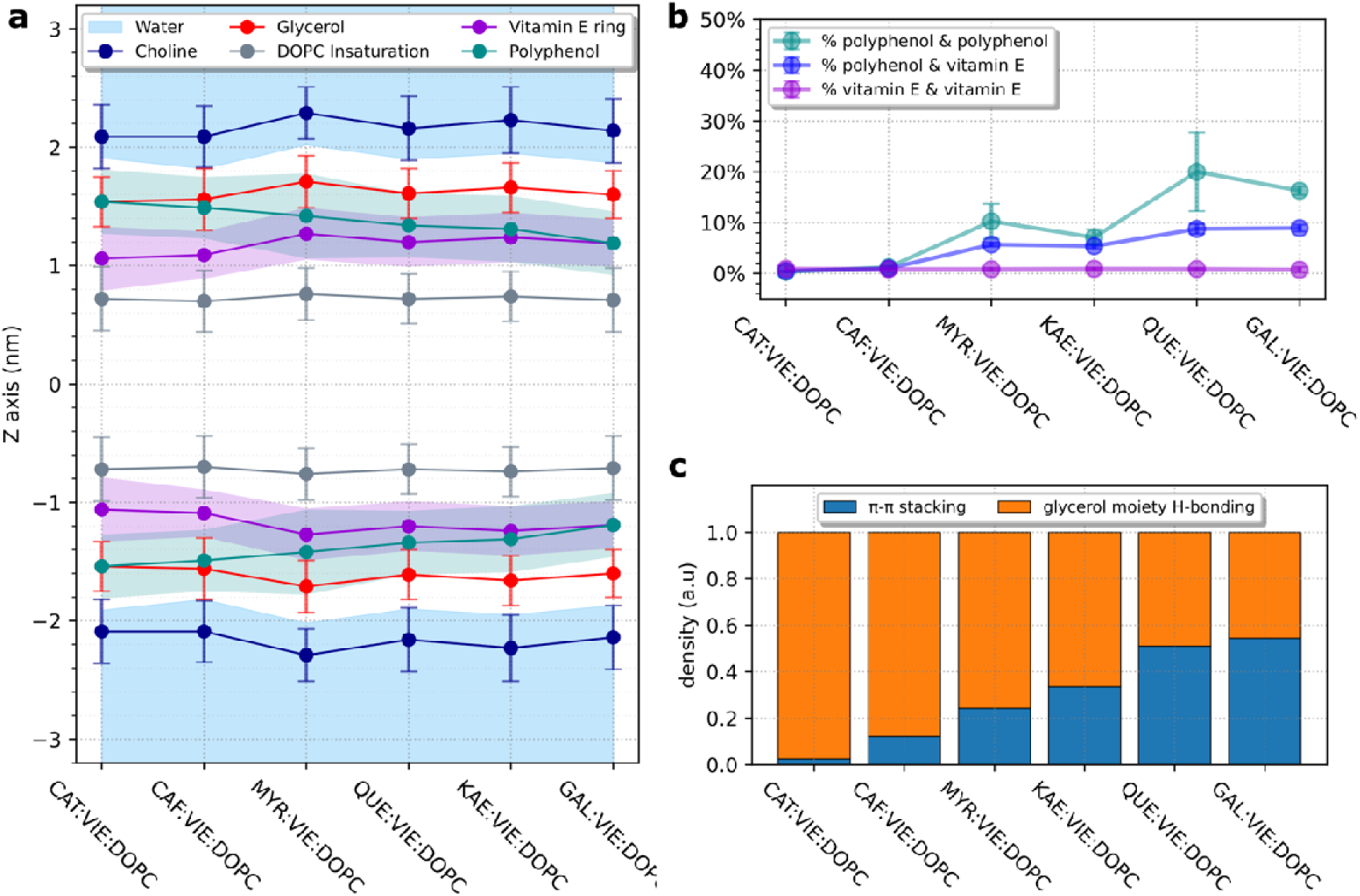
(a) Time-averaged position occupied by lipid moieties, the six polyphenols and water at 50% of the electron profile density along z-axis; (b) percentage of π-π stacking interaction between the two partners along the MD simulations in DOPC bilayer membrane; (c) normalized ratio distribution of polyphenol-vitamin E (VIE) π-π stacking interaction and H-bonding between the DOPC glycerol moieties and the polyphenols, when the molecules are in the sphere-of-action. The six polyphenols are: catechin (CAT), caffeic acid (CAF), myricetin (MYR), kaempferol (KAE), quercetin (QUE), galangin (GAL)

Interestingly, the same ranking was observed when assessing the radical scavenging capacity of lipid peroxidation in liver microsomes^56^ for these six compounds. The antioxidant activity in the presence of a small amount of vitamin E was ranked as follows: galangin > quercetin > kaempferol > myricetin > catechin. This means that increasing the lipophilicity (*i.e.* by removing hydroxyl group from the flavonoid B-ring) does not linearly lead to an improved antioxidant capacity or a better synergistic effect with vitamin E. Instead, the location of the polyphenol-regenerator appears as a much instructive complementary descriptor, which can be efficiently assessed by MD simulations.

## 3 Conclusion

The goal of this work was to benchmark classical all-atom MD simulations for their capacity to reproduce dipole-dipole interactions, and to gain further insight into the observed fluorescence quenching data obtained for two prototypical compounds (quercetin and vitamin E) in DOPC lipid bilayers. For this purpose, the molecular systems were chosen to mimic the corresponding experimental data, available from the literature. The thorough analysis of MD trajectories confirmed a very good agreement between theoretical and fluorescence quenching data. A sphere-of-action model was considered, in which a combined static quenching is likely to occur, as fitted in the modified Stern-Volmer plot. The contribution of static quenching (∼10%) was attributed to the formation of π-π stacking complexes, which was predicted by MD simulations with high accuracy compared to experimental values. Both polyphenol self-interaction and polyphenol-vitamin E hetero-association are likely to occur. The former mechanism is predominant when polyphenol concentration increases, leading to the formation of larger polyphenol aggregates, which are precursors of crystallization related to toxicity in the human body ^57^. Conversely, the latter interactions (π-π stacking between polyphenols with vitamin E) is more likely associated with beneficial effects on the organism, as it seems highly correlated to the capacity of antioxidant activity and regeneration. In other words, MD simulations are proven reliable at capturing, describing, and quantifying the underlying processes leading to synergism within π-π stacking complexes involving two different π-conjugated antioxidants. Here we highlight that for industrial application and the search for the best antioxidant cocktails, MD simulations can support the prediction of non-covalent association in different environments, following both the amount of π-π stacking but also a more global descriptor, namely the amount of molecular contacts with a sphere of action. An accurate description of these two atomistic numbers has proved crucial for a very practical use in the development of powerful antioxidant cocktails.

## 4 Abbreviations

BDE: Bond Dissociation Enthalpy
DOPC: 1,2-dioleoyl-sn-glycero-3-phosphocholine
HAT: Hydrogen Atom Transfer
MD: Molecular Dynamics
ROS: Reactive Oxygen Species
SV: Stern-Volmer

## 5 Supporting Information

Figure S1. Percentage of π-π stacking interaction when the starting configuration of the molecules is random or pre-stacked.

Figure S2. Vitamin E emission spectrum at (291nm) and quercetin absorption spectrum overlap.

Figure S3. Difference of charges obtain from REStrain electrostatic Potential (RESP) with Duan et. al. model in the 4 different flavonoids represented in shade of color.

Table S2. Ratio data of vitamin E and quercetin simulations.

Table S3. Ratio data of vitamin E and polyphenols simulations.

## 6 Acknowledgments

We thank CALI (CAlcul en Limousin), Florent Di Meo and Xavier Montagutelli for computational resources as well as InSiliBio and INSERM for financial support. The work in Madrid was supported by the Spanish Ministry for Science through the MICIN-FEDER project PID2022-138222NB-C21 and though the Severo Ochoa program for Centers of Excellence in R&D (CEX2020-001039-S).

## Footnotes

‡ It is possible to distinguish between dynamic and static quenching by measuring the fluorescence lifetime in presence (absence) of the quencher, τ_F_ (τ_F,0_). For exclusive collisional quenching, τ_F,0_/τ_F_ = Φ_F,0_/Φ_F_, while for exclusive static quenching τ_F,0_/τ_F_ = const; see Ref. ^36^ Unfortunately, time-resolved fluorescence data were not reported in Ref. ^16^

§ The quadratic model of eq. (3) gives somewhat different values of K_sv_ = 3.68 M^-1^, and V_m_ = 0.93 M^-1^, indicating that Vm is not sufficiently small to allow for a reliable replacement of eq. (2) by eq. (3)

## Notes

### Competing Interest Statement

The authors have declared no competing interest.

