## Supporting information for "Predicting Non-Covalent Interactions Between Antioxidants In Biological Membranes Through Molecular Dynamics"

a) INSERM U1248 Pharmacology & Transplantation, Univ. Limoges, CBRS, 2 rue du prof. Descottes, F-87000 Limoges, France

b) InSiliBio, 1 avenue d’Ester, Ester Technopôle, F-87000 Limoges, France

c) Madrid Institute for Advanced Studies, IMDEA Nanoscience, Ciudad Universitaria de Cantoblanco, Calle Faraday 9, 28049 Madrid, Spain

d) RCPTM CATRIN Palacky University, Slechtitelu 27, 783 71, Olomouc, Czech Republic.

### 1) Assessing vitamin E and quercetin concentration within the liposome.

In the supplementary data of Fabre *et al.* paper^1^, the concentration of vitamin E and quercetin were given within the suspension as 50µM and between 0 and 100µM, respectively. Vitamin E is a lipid-soluble molecule and so entirely inserts into the DOPC liposomes. Also, according to the literature quercetin has a partition coefficient in DOPC of *ca.* 8.41 ± 1.51*10^4^, representing approximately 99% of the molecules inserted in the lipid compartment of the liposome^2^.

Therefore, we can easily assume that the two molecules are essentially inserted in the formed liposomes. However, their respective concentration is expected to be higher than in the suspension because the liposome volume is smaller. Nevertheless, we need the actual quercetin and vitamin E concentration within the liposome to correctly assess the Stern-Volmer constant of fluorescence quenching. The calculation of these concentrations is given as follows. The formed liposomes are likely large unilamellar vesicles (LUV) because of the reported diameter (159 nm) and the preparation process within which [vitamin E] = 4 x [DOPC].

In our simulation, we used **72 lipid molecules** organized as a bilayer covering a surface area of **2.5**⋅**10^-15^ dm²** (coming from **5 nm x 5 nm, the size of the simulation cubic box**). This results in a surface area per (DOPC) lipid of **3.5 x 10^-17^ dm²**. To calculate the total number of DOPC molecules in a liposome with a diameter of **159 nm**, we first determined the surface area of the liposome, which is approximately **7.94**⋅**10^-12^ dm² (4**⋅**π**⋅**r^2^)**. Using the surface area per bilayer, we can estimate that a liposome of this size contains about **228737 DOPC molecules**.

To find the concentration of DOPC in the liposomes, we need to divide the total number of DOPC molecules by the volume of the liposome bilayer. Assuming a bilayer thickness of 5 nm, we can calculate the volume of the spherical cap by:

$$Volume = \frac{4}{3}\cdot\pi\cdot(r^{3}-{r'}^{3})$$

Where: r (= 159nm/2) is the outer radius of the liposome, and r′ ( = r-thickness = 154nm/2) is the inner radius. This gives a volume of approximately 1.92⋅10^-19^ L.

Using the Avogadro's number (6.023⋅10^23^ mol^-1^) and the liposome volume, the number of DOPC molecules is converted into 2 M concentration.

Since the concentration of vitamin E is one-fourth of DOPC, we can deduce that:

[Vitamin E] = [DOPC]/4 = 0.5 M. Likewise, the concentration of quercetin varies from **0 to 1 M according to the ratio mentioned in the article**.

2) **Comparing starting configuration of simulation.**

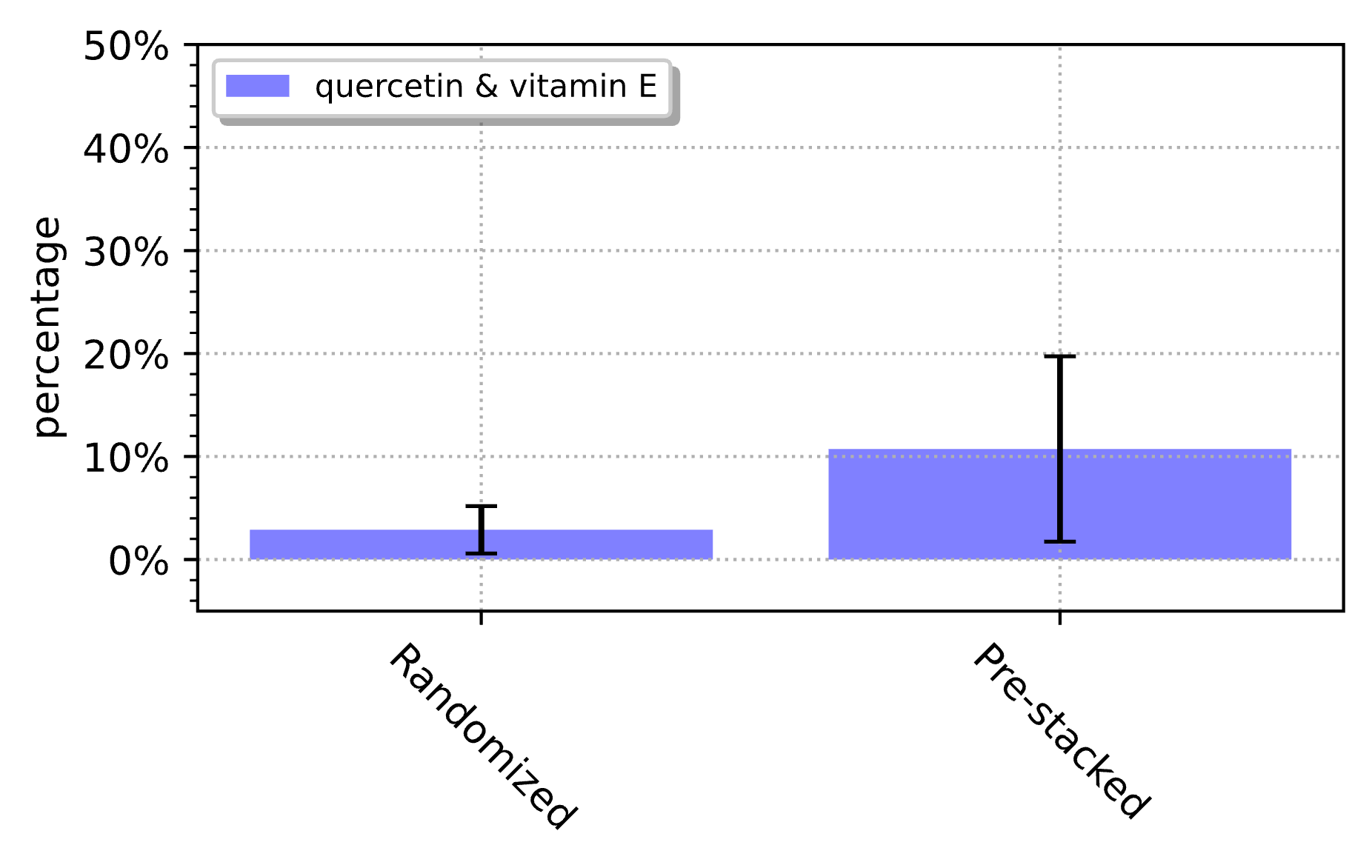

**Figure S1. Percentage of stacking interaction when the starting configuration of the molecules is random or pre-stacked.**

Percentage are counted within system simulation (not within sphere-of-action) of quercetin:vitaminE:DOPC = 4:4:128. Ring center distance was set to 5Å and the angle between the two orthogonal vectors to the ring was set to 20°.

3) **Reabsorption: Spectrum overlap and event probability**

Fluorescence quenching in the membrane can be caused by different physical mechanisms such as photoinduced electron transfer or resonant energy transfer, however by 'trivial' radiative transfer, i.e. reabsorption. Here, with both a significant overlap between the spectrum of emission of the fluorophore and the spectrum of absorption of the quencher, and a sufficiently high concentration of quencher, the fluorescence can be absorbed by the surrounding quencher molecules. In the case of vitamin E and quercetin, the first condition is met as the respective emission and absorption spectra overlap between *c.a.* 300-400 nm (see below). The second condition is likely to be fulfilled as the concentration of quencher inside the lipid bilayer is high with C_Q_ of up to 0.09 M (Figure 1). The probability 𝜑 for the reabsorption of vitamin E fluorescence by quercetin is calculated by equation (2)^3,4^**.**

|  | 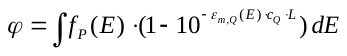 | (2) |
| --- | --- | --- |

where *f_P_*(*E*) is the area-normalized fluorescence spectrum of the probe fluorophore ***P***, ε_m,Q_(*E*) is the absorption spectrum of ***Q*** in units of molar extinction coefficients, and *L* is the path length. Hence, instead of Φ*_F_*, an apparent quantum yield Φ*_F_*' is observed.^5^

$$\Phi_{F}'= \frac{(1- \varphi)\cdot\Phi_{F}}{1-\varphi\cdot\Phi_{F}}$$

(3)

Quenching by an encounter complex in the presence of reabsorption is given by a non-linear Stern-Volmer (SV) equation:^6^

$$\frac{I_{0}}{I}= \frac{1+ K_{sv}\cdot C_{Q} - \varphi\cdot\Phi_{F}}{1-\varphi}$$

(4)

where C_Q_ represents the concentration of quercetin, 𝜀_m_ is the extinction coefficient, *d* is the light path length (1cm usually) and Φ_F_ is the fluorescence quantum yield. It is noted that in the absence of quenching (i.e. *K_SV_* = 0), eq. (4) indeed reduces to eq. (3), while in the absence of reabsorption (ϕ = 0), it reduces to the simple *Stern-Volmer* equation.

When fitting the experimental data to this equation, 𝜑 was very close to 1 making the resulting I_o_/I values irrelevant (see below).

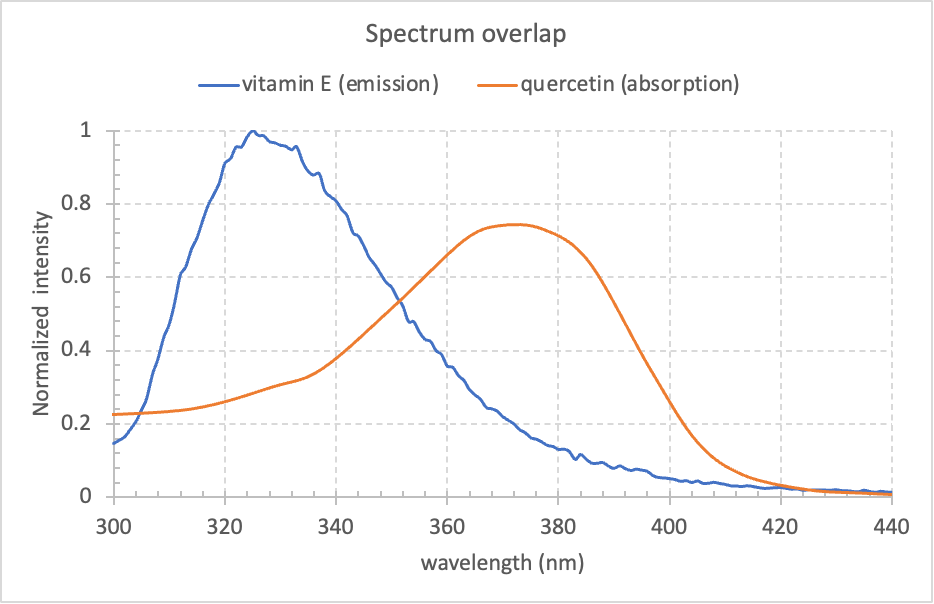

**Figure S2. Vitamin E emission spectrum at (291nm) and quercetin absorption spectrum overlap.**

Quercetin absorption spectrum was retrieved from the spectrabase web database.

**Table S1.** Probability of secondary inner-filter effect (reabsorption)

| Extinction coefficient (ε_m_) | [Quercetin] (C_Q_) | Probability (ϕ) | K_sv_ | Io/I |
| --- | --- | --- | --- | --- |
| = 8158 M^-1^ cm^-1^  *For d=1cm, from* ^6^ | 0.1 M | 1 | 1 | Inf |
|  | 0.2 M | 1 | 1 | Inf |
|  | 0.25 M | 1 | 1 | Inf |
|  | 0.3 M | 1 | 1 | Inf |
|  | 0.4 M | 1 | 1 | Inf |
|  | 0.5 M | 1 | 1 | Inf |
|  | 0.7 M | 1 | 1 | Inf |
|  | 0.8 M | 1 | 1 | Inf |
|  | 1 M | 1 | 1 | Inf |

### 4) Raw ratio data of molecule pair, in sphere-of-action, in π-π stacking and their corresponding radius

**Table S2. Ratio data of vitamin E and quercetin simulations.** Vitamin E (**V**) and quercetin (**Q**); concentration of quercetin (**[Q]_lip_**) and vitamin E (**[V]_lip_**) in liposome (**lip**); **I_o_/I** is the ratio of fluorescence intensity; **Rad** is the radius of the two molecules sphere-of-action considered; $\boldsymbol{\%R}_{\boldsymbol{[Q]}}^{\boldsymbol{in sphere}}$ and $\boldsymbol{\%R}_{\boldsymbol{[Q]}}^{\boldsymbol{\pi-\pi stacking}}$ are respectively the percentage of molecules contained within the sphere-of-action considered along MD simulation and the percentage of π-π stacked molecules within the sphere-of-action. Ratio was assessed for each system of concentration ([Q] in subscript).

| [Q]_lip_  (mM) | [V]_lip_  (mM) | Ratio  ([Q]/[V])_lip_ | Io/I | QV  Rad (Å) | $\boldsymbol{\%R}_{\boldsymbol{[Q]}}^{\boldsymbol{QV in sphere}}$ | $\boldsymbol{\%R}_{\boldsymbol{[Q]}}^{\boldsymbol{QV \pi-\pi stacking}}$ | QQ  Rad (Å) | $\boldsymbol{\%R}_{\boldsymbol{[Q]}}^{\boldsymbol{QQ in sphere}}$ | $\boldsymbol{\%R}_{\boldsymbol{[Q]}}^{\boldsymbol{QQ \pi-\pi stacking}}$ | VV  Rad (Å) | $\boldsymbol{\%R}_{\boldsymbol{[Q]}}^{\boldsymbol{VV in sphere}}$ | $\boldsymbol{\%R}_{\boldsymbol{[Q]}}^{\boldsymbol{VV \pi-\pi stacking}}$ |
| --- | --- | --- | --- | --- | --- | --- | --- | --- | --- | --- | --- | --- |
| 0 | 250 | 0 | 1 | 7 | 0 | 0 | 8 | 0 | 0 | 6 | 0 | 0 |
| 100 | 250 | 0.2 | 1.5 |  | 30.40 ± 3.95 | 8.73 ± 1.81 |  | 12.07 ± 7.17 | 11.97 ± 16.78 |  | 13.27 ± 0.75 | 0.80 ± 0.00 |
| 200 | 250 | 0.4 | 2.05 |  | 44.42 ± 0.66 | 9.40 ± 1.37 |  | 30.38 ± 3.20 | 9.77 ± 11.57 |  | 14.72 ± 1.35 | 1.23 ± 0.29 |
| 250 | 250 | 0.5 | 2.36 |  | 53.88 ± 2.67 | 9.83 ± 0.62 |  | 40.48 ± 7.50 | 12.53 ± 5.86 |  | 12.51 ± 1.03 | 0.87 ± 0.12 |
| 300 | 250 | 0.6 | 2.72 |  | 54.79 ± 5.50 | 6.87 ± 0.95 |  | 39.09 ± 2.77 | 15.43 ± 5.53 |  | 14.28 ± 0.84 | 0.77 ± 0.25 |
| 400 | 250 | 0.8 | 3.38 |  | 65.06 ± 5.20 | 7.67 ± 0.59 |  | 36.75 ± 2.24 | 19.40 ± 3.26 |  | 11.99 ± 0.96 | 0.90 ± 0.08 |
| 500 | 250 | 1 | 4.16 |  | 71.72 ± 3.76 | 8.77 ± 0.96 |  | 46.77 ± 2.57 | 20.00 ± 7.72 |  | 11.16 ± 0.55 | 0.87 ± 0.19 |
| 700 | 250 | 1.4 | 5.9 |  | 81.35 ± 1.24 | 7.63 ± 0.33 |  | 71.09 ± 3.04 | 19.73 ± 2.64 |  | 12.34 ± 0.38 | 0.90 ± 0.28 |
| 800 | 250 | 1.6 | 6.88 |  | 86.89 ± 2.06 | 8.03 ± 0.42 |  | 86.24 ± 5.19 | 16.83 ± 3.03 |  | 10.18 ± 1.84 | 0.77 ± 0.24 |
| 1000 | 250 | 2 | 9.03 |  | 87.61 ± 3.14 | 6.63 ± 1.07 |  | 100.29 ± 2.12 | 19.70 ± 0.57 |  | 10.37 ± 0.47 | 1.27 ± 0.41 |

**Table S3. Ratio data of vitamin E and polyphenols simulations.** Each number of polyphenols (**P**) and vitamin E (**V**) molecules in molecular dynamic simulation (**MD**); **Rad** is the radius of the two molecules sphere-of-action considered; $\boldsymbol{\%R}_{\boldsymbol{[P]}}^{\boldsymbol{in sphere}}$ and $\boldsymbol{\%R}_{\boldsymbol{[P]}}^{\boldsymbol{\pi-\pi stacking}}$ are respectively the percentage of molecules contained within the sphere-of-action considered along MD simulation and the percentage of π-π stacked molecules within the sphere-of-action. Ratio was assessed for each system of concentration ([P] in subscript).

| Polyphenols (P) | ^8^AOX power ranking | Ratio (P/V)_MD_ | PV  Rad (Å) | $\boldsymbol{\%R}_{\boldsymbol{[P]}}^{\boldsymbol{PV in sphere}}$ | $\boldsymbol{\%R}_{\boldsymbol{[P]}}^{\boldsymbol{PV \pi-\pi stacking}}$ | PP  Rad (Å) | $\boldsymbol{\%R}_{\boldsymbol{[P]}}^{\boldsymbol{PP in sphere}}$ | $\boldsymbol{\%R}_{\boldsymbol{[P]}}^{\boldsymbol{PP \pi-\pi stacking}}$ | VV  Rad (Å) | $\boldsymbol{\%R}_{\boldsymbol{[P]}}^{\boldsymbol{VV in sphere}}$ | $\boldsymbol{\%R}_{\boldsymbol{[P]}}^{\boldsymbol{VV \pi-\pi stacking}}$ |
| --- | --- | --- | --- | --- | --- | --- | --- | --- | --- | --- | --- |
| Catechin | 6 | 1 | 7 | 61.35 ± 0.81 | 0.47 ± 0.09 | 7 | 37.82 ± 1.17 | 0.33 ± 0.09 | 6 | 16.02 ± 0.26 | 0.73 ± 0.65 |
| Caffeic acid | 5 |  | 6.5 | 47.50 ± 3.88 | 0.97 ± 0.05 | 8 | 28.03 ± 1.96 | 1.20 ± 0.33 |  | 14.65 ± 2.13 | 0.83 ± 0.69 |
| Myricetin | 4 |  | 7 | 67.96 ± 1.90 | 5.70 ± 0.45 | 8 | 37.39 ± 0.64 | 10.27 ± 3.43 |  | 16.56 ± 2.17 | 0.83 ± 0.25 |
| Kaempferol | 3 |  | 7 | 72.37 ± 0.36 | 5.37 ± 0.52 | 8 | 40.65 ± 0.84 | 7.10 ± 1.59 |  | 12.50 ± 2.58 | 0.90 ± 0.22 |
| Quercetin | 2 |  | 7 | 71.72 ± 3.76 | 8.77 ± 0.96 | 8 | 46.77 ± 2.57 | 20.00 ± 7.72 |  | 11.16 ± 0.55 | 0.87 ± 0.19 |
| Galangin | 1 |  | 7 | 75.05 ± 4.29 | 8.93 ± 0.66 | 8 | 60.18 ± 2.25 | 16.27 ± 0.84 |  | 12.65 ± 1.15 | 0.73 ± 0.54 |

### 5) Comparing atomic partial charges of flavonoid

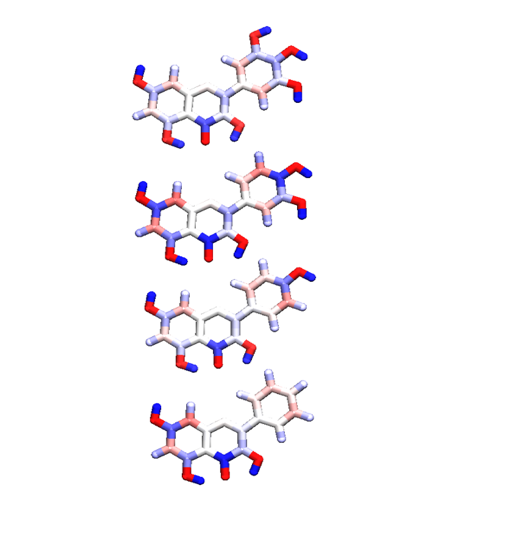

quercetin

galangin

kaempferol

myricetin

**Figure S3.** **Difference of charges obtain from REStrain electrostatic Potential (RESP^9^) with Duan et. al.^10^ model in the 4 different flavonoids represented in shade of color.**

Redish for negative charges and blueish for positive charges
